# Exploring vulnerable building blocks in protein-protein interaction networks of breast tumor and adjacent normal tissues

**DOI:** 10.1101/2025.02.26.640272

**Authors:** Swapnil Kumar, Avantika Agarwal, Vaibhav Vindal

**Affiliations:** Department of Biotechnology & Bioinformatics, School of Life Sciences, University of Hyderabad, Gachibowli, Hyderabad, 500046, India

**Keywords:** Tumor-adjacent normal tissues, breast cancer subtypes, vulnerable proteins, protein complex, network vulnerability analysis, influence analysis, escape velocity centrality

## Abstract

Tumor-adjacent normal tissues (TANTs) histologically and morphologically look normal and are commonly used as a control in patient-based cancer studies. Previous studies have revealed that TANTs present a unique transitional state between healthy normal and tumor tissues. However, there is little or no knowledge about the landscape of protein-protein interactions (PPIs) in TANTs and how they differ from the tumor tissues. Here, we integrate the PPI data mapped onto the differentially expressed genes in TANTs and tumor tissues compared to healthy normal tissues to comprehensively construct and analyze the PPI networks of TANTs and breast tumor tissues (viz., Luminal A, Luminal B, Her2, Basal, and Normal-Like). First, these PPI networks were analyzed using network influence and vulnerability analyses from the NetVA R package. Consequently, our study revealed 134 vulnerable proteins (VPs), 21 vulnerable protein pairs (VPPs), and 94 influential proteins (IPs) commonly shared across all six tissue networks. Further, we identified a set of 34 proteins as common hubs and another set of seven proteins as common bottlenecks across all six tissue networks. Next, these VPs, IPs, hubs, and bottlenecks were investigated for their associations with various diseases, including cancers, and found to share a significant number of well-known cancer-associated proteins, viz., AR, BRCA1, ERBB2, FN1, FOXA1, JUN, MKI67, and NRAS. Thus, by applying network vulnerability, influence, and gene-disease association-based analyses, we suggest lists of known and novel candidates along with their associated protein complexes potentially involved in breast cancer tumorigenesis and present across TANTs and different breast cancer subtypes.

## Introduction

Proteins perform collectively by interacting with other proteins and forming complexes for their molecular functions, processes, and pathways [Alberts, 1998; Gavin et al., 2006]. The interaction of proteins with each other leads to the formation of network-like structures. These networks are composed of proteins as nodes and interactions among them as edges, and these can be of various types and can represent a disease-, process-, or even a normal system. Any perturbation in a network representing a normal state, such as the insertion, deletion, or inactivation of nodes, results in a dysregulated or perturbed system with altered functions and processes. This perturbed or dysregulated network leads to the development of a disease [Rai et al., 2014; Jalan et al., 2015]. The protein-protein interaction (PPI) networks of tumor conditions in the case of breast and cervical cancers were observed to be perturbed compared to the normal ones. This indicated that a perturbation in the normal system led to the development of a disease system.

Breast cancer causes high morbidity and mortality among women and is the most commonly diagnosed cancer type across the globe. It is a heterogeneous disease encompassing five different subtypes based on hormone receptors: ER (Estrogen receptor), PgR (Progesterone receptor), and HER2 (Human epidermal growth factor receptor 2). These molecular subtypes are Luminal A (LumA) (ER+/PgR+, HER2-, and Ki67 low), Luminal B (LumB) (ER+/PgR+, HER2- or HER2+, and Ki67 high), HER2-enriched (Her2) (ER+/PgR+ and HER2+), Basal-Like (Basal) (ER-/PgR- and HER2-), and Normal-Like (NormL) (ER+/PgR+, HER2-, and Ki67 low) [Dai et al. 2015]. The current treatment regimen for this disease is not able to manage and treat it efficiently due to various reasons such as disease recurrence, advanced stage, distant metastasis, and chemoresistance. Further, breast cancer or any cancer studies utilize the tumor tissues as treated samples and the tumor-adjacent normal tissues (TANTs) as control samples to understand the tumorigenesis and progression of the disease, which further helps to identify and develop diagnostic and therapeutic strategies. However, the TANTs appear to be morphologically normal but are molecularly altered and have various tumorigenic features, including altered gene expression and mutation profiles, compared to the completely normal, i.e., healthy tissues [Aran et al., 2017; Kumar & Vindal, 2023]. Therefore, the TANTs need to be re-explored along with tumor tissues to better understand the disease development and progression towards developing more efficient and novel diagnostics and therapeutics, especially for the early stage of cancer.

Detection of key molecular players of a disease such as breast cancer forms a decisive part of an analysis of a disease-specific biological network, leading to a better understanding of that disease and, hence, the generation of potential methods of diagnostics and therapeutics. Computational construction and analysis of network models representing cancers have previously been utilized for a detailed understanding of underlying pathways and related mechanisms [Srivastava et al., 2014; Rai et al., 2014; Jalan et al., 2015; Li et al., 2020; Tiwari & Dwivedi, 2019]. There are various network analysis approaches, and network vulnerability analysis (NVA) is one of the most important among them, as evident from previous studies [Abdi et al., 2008; Dartnell et al., 2005; Podder et al., 2018]. The NVA assesses significant associations of a network’s functional essentiality with its topological properties [Kumar et al., 2024]. Besides, some node centralities, like escape velocity-based centrality (EVC) and its extended version (EVC+), are also used to detect key nodes that outperform other topological properties of nodes, viz., degree and betweenness [Ullah et al., 2022; Kumar et al., 2024].

Moreover, protein complexes represent groups of proteins that are densely connected within their respective groups but are sparsely connected with proteins of other groups or components of the network. Recent advancements in experimental approaches, especially the integration of affinity purification with mass spectrometry (AP-MS), have produced an enormous amount of PPI data for several organisms, including humans. These PPI data of humans are publicly available in several databases that can be utilized for the construction of PPI networks of TANTs and breast cancer subtypes. Further, these networks can be analyzed for protein complex detection, which is crucial to understanding cellular organization, behavior, and functions.

In the current study, six PPI networks (five subtypes and one TANT) were constructed using lists of DEGs of different subtypes and TANTs of breast tumors as retrieved from our previous study coupled with PPIs collected from various sources. Subsequently, these networks were analyzed using the approach of NVA to identify key vulnerable proteins (VPs) and protein pairs (VPPs). Additionally, these VPs were used to predict vulnerable protein complexes (VPCs) in the case of all six tissue networks. These VPs, VPPs, and VPCs can be explored further to better understand the early stage-specific mechanisms of tumor development of different subtypes of breast cancer through TANTs and design better therapeutic and diagnostic candidates.

## Methods

### Construction of protein interaction networks

Differentially expressed genes (DEGs) for each tissue type, viz., tumors (Basal, Her2, LumA, LumB, and NormL) and TANTs, were collected from our previously published work [Kumar & Vindal, 2023]. These DEGs were mapped to PPIs from four databases: HuRI [Luck et al., 2020], HIPPIE [Alanis-Lobato et al., 2017], CancerNet [Meng et al., 2015], and PIPs [McDowall et al., 2009]. Consequently, six PPI networks were constructed, one for each tissue type, including five subtypes and one TANT. A PPI network may contain only a single, completely connected component or one large component along with one or more than one disconnected smaller component. In the present study, only the largest connected component of each network was considered for further analysis.

### Identification of hubs and bottlenecks

First, all six PPI networks (five subtypes and one TANT) were analyzed to identify hubs and bottlenecks for each tissue network. A protein with a high degree value is considered a hub, whereas a protein with a high betweenness value is considered a bottleneck. The “detectHubs” and “detectBottlenecks” functions of the NetVA R package [Kumar et al., 2024] were used to identify hubs and bottlenecks, respectively. These functions use the Pareto principle of 80:20 to identify hubs and bottlenecks. The human PPIs can have false positive interactions, which may result in biased calculation of degree and betweenness values. To reduce the chance of this bias, 5% of all edges in the PPI network were shuffled 20 times, each time with different sets of edges, resulting in the construction of 20 rewired networks. For each of these 20 rewired networks, hubs and bottlenecks were identified. Thus, all proteins present as hubs in all these 20 networks were only considered as hubs. Similarly, all proteins present as bottlenecks in all these 20 networks were only considered as bottlenecks.

Degree: It is defined for a node as the number of neighboring nodes directly connected with that node in a given network.

Betweenness: It is defined for a node as the ratio of all the shortest paths that pass through that node and all the shortest paths of every node pair present in the given network.

### Network vulnerability analysis

The largest connected components of all six PPI networks were analyzed one by one using the NVA approach, which is available in the NetVA R package. The “netva” function of the NetVA package implements vulnerability analysis utilizing 14 different topological properties. For this, first, all proteins participating in the PPI network were removed one by one, and the values of these 14 topological properties were calculated for each resultant network after removing a node. For all 14 properties, the most vulnerable proteins (VPs) were identified using the outlier detection criteria implemented in a boxplot. Second, all possible protein pairs of these VPs (where two VPs interact with each other in the PPI network) were removed from the original network, and again, those 14 topological properties were calculated for the resultant network after the removal of the protein pairs. Subsequently, the most vulnerable protein pairs (VPPs) were identified using the same procedure as VPs. The topological properties implemented in the NetVA package are articulation point number, average closeness centrality, average betweenness centrality, average eccentricity, average node connectivity, clustering coefficient, average path length, heterogeneity, network diameter, cohesiveness, compactness, global efficiency, network density, and network centralization.

Articulation point: It represents a node or protein whose deletion or removal from its network results in two or more disconnected components of that network, i.e., sub-networks. The number of disconnected subnetworks of a given network after removing a node is referred to as the articulation point number of that node.

Average betweenness centrality: It is the average value of the betweenness centrality of all nodes present in a given network.

Average closeness centrality: It is the average value of closeness centrality (CC) of all nodes present in a given network, and the CC of a node is defined as the inverse of the sum of distances of that node from all the other nodes present in the given network [Freeman, 1978].

Average eccentricity: It is the average value of eccentricity centrality (EC) of all nodes present in a network. The EC of a node can be defined as the inverse of the longest of all shortest paths of that node from all the other nodes present in a network [Hage & Harary, 1995].

Average node connectivity: It is the average degree value of all the nodes present in a given network.

Average path length: It is the average value of all the shortest paths among all the node pairs present in a given network.

Clustering coefficient: It is defined as the ratio of the number of edges between the neighbors of a node and the number of all edges that are possible between all those neighbors of that node present in a given network.

Cohesiveness: It is the measurement of the likeliness of a group of proteins that can form a protein complex [Nepusz et al., 2012], and hence it is defined as the ratio of the edge weight of proteins present in a group to the sum of edge weight of proteins of that group and edge weight of proteins connecting the group with other parts of the network.

Compactness: It is the ratio of the number of cliques of three nodes in a subnetwork to the total number of cliques of three nodes present in that subnetwork and at the boundary of the subnetwork, which connects the subnetwork with other parts of the network [Zhang & Zou, 2015].

Global efficiency: It is the average value of the inverse of distances between all node pairs in a network [Vragović et al., 2005; Latora & Marchiori, 2001].

Heterogeneity: It is defined as the ratio of the standard deviation to the average value of the degree of all nodes present in a given network [Dong & Horvath, 2007].

Network centralization: It is defined as the basic index of a network that is widely used to check connectivity distribution therein [Freeman, 1978].

Network density: It is the ratio of the number of all actual edges and the number of all feasible edges in a network [Wasserman & Faust, 1994].

Network diameter: It is defined as the longest path length out of all shortest path lengths between every node and all other nodes present in a given network.

### Network influence analysis

Subtypes and TANT networks were also analyzed using escape velocity centrality (EVC), which revealed each network’s most influential proteins (IPs). The “evc” function of the NetVA R package was used for this. The “evc” function calculates the values of EVC and EVC+ (extended version of EVC) for each protein present in the network, and proteins commonly present among the top 20 percentile of all the proteins based on their EVC and EVC+ values were considered the IPs.

The EVC and EVC+ of a node ‘i’ can be defined as follows:

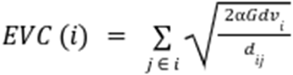

where α is a tunable factor with a value between 0.1 and 1, which controls the degree effect; G is the constant with a value of 1; dvi is the degree of node ‘i’, dij is the shortest path length using Dijkstra’s algorithm between nodes ‘i’ and ‘j’; jєi is the set representing neighboring nodes of node ‘i’.

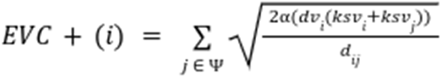

where ksvi and ksvj are the values of k-shell of nodes ‘i’ and ‘j,’ respectively, and jɛψ is the set representing neighboring nodes. The k-shell value of all nodes of a maximal subnetwork is k, where the maximal subnetwork is composed of nodes with degrees at least k [Ullah et al., 2022].

### Identification of vulnerable protein complex

Proteins do not work individually but rather work by interacting with each other and forming protein complexes. These protein complexes play vital roles in the execution of various cellular pathways, molecular functions, and biological processes. All the protein complexes formed by VPs were identified for each network using ClusterONE v1.0, an open-source software [Nepusz et al. 2012]. ClusterONE (Clustering with Overlapping Neighborhood Expansion) implements a network or graph clustering algorithm that handles both weighted and non-weighted graphs and quickly detects overlapping clusters within the graph. It is useful, especially for detecting protein complexes with confidence values in PPI networks. To detect vulnerable protein complexes (VPCs), all possible protein complexes with a minimum of three participating proteins were first predicted in each tissue type. Further, previously identified VPs of each tissue were mapped onto the identified protein complexes of the same tissue to identify corresponding VPCs.

### Association of diseases with key proteins

All identified key proteins in this study, viz., VPs, IPs, hubs, and bottlenecks, were checked for their association with any diseases, including cancers, using the DisGeNET database [Piñero et al., 2020]. The DisGeNET integrates curated and text-mined information on associations of genes and variants with human diseases derived from various specialized repositories in different disease categories. Only gene-disease associations reported in more than one published literature were considered to identify the significant associations of key proteins with diseases.

## Results

### Protein-protein interaction networks of tumor tissues and TANTs

PPI networks of tumor tissues and TANTs were constructed by collecting edges as discussed in the Methods section, and the detailed distribution is given in Table 1.

**Table 1.** Distribution of nodes and edges across all tissue types.

| **Network** | **DEGs** | **Nodes** | **Edges** |
| --- | --- | --- | --- |
| Basal | 7846 | 7190 | 83640 |
| Her2 | 8344 | 7634 | 90669 |
| LumA | 7175 | 6472 | 61303 |
| LumB | 7917 | 7266 | 85536 |
| NormL | 6090 | 5317 | 41221 |
| TANT | 4313 | 3526 | 18505 |

### Hubs and bottlenecks of tumor and TANT networks

Each tissue network was first analyzed to detect hubs and bottlenecks using the “detectHubs” and “detectBottlenecks” functions of the NetVA package. These identified hubs and bottlenecks had a very little or negligible chance of any false positive because of the inclusion of 5% rewired interactions as a negation factor implemented in these NetVA functions against any false positive interactions in the human PPI data. The distributions of hubs, bottlenecks, and hubs as well as bottlenecks are available in Figs. 1, 2, and 3. The detailed lists of hubs and bottlenecks across all six tissues are available in Tables S1 and S2.

**Figure 1.**
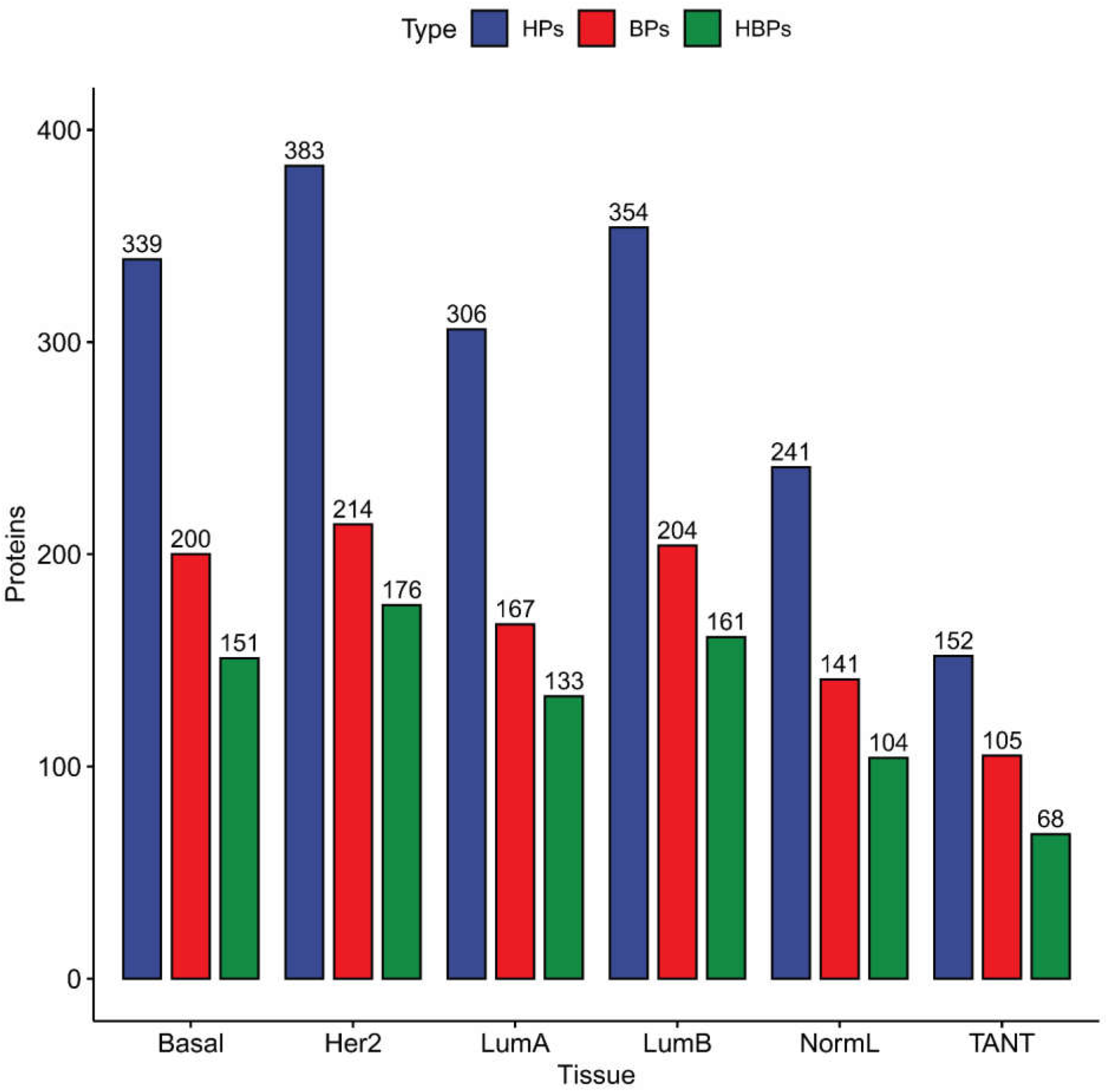
Distribution of proteins which are hubs (HPs), bottlenecks (BPs), and hubs as well as bottlenecks (HBPs) across all six tissue networks

**Figure 2.**
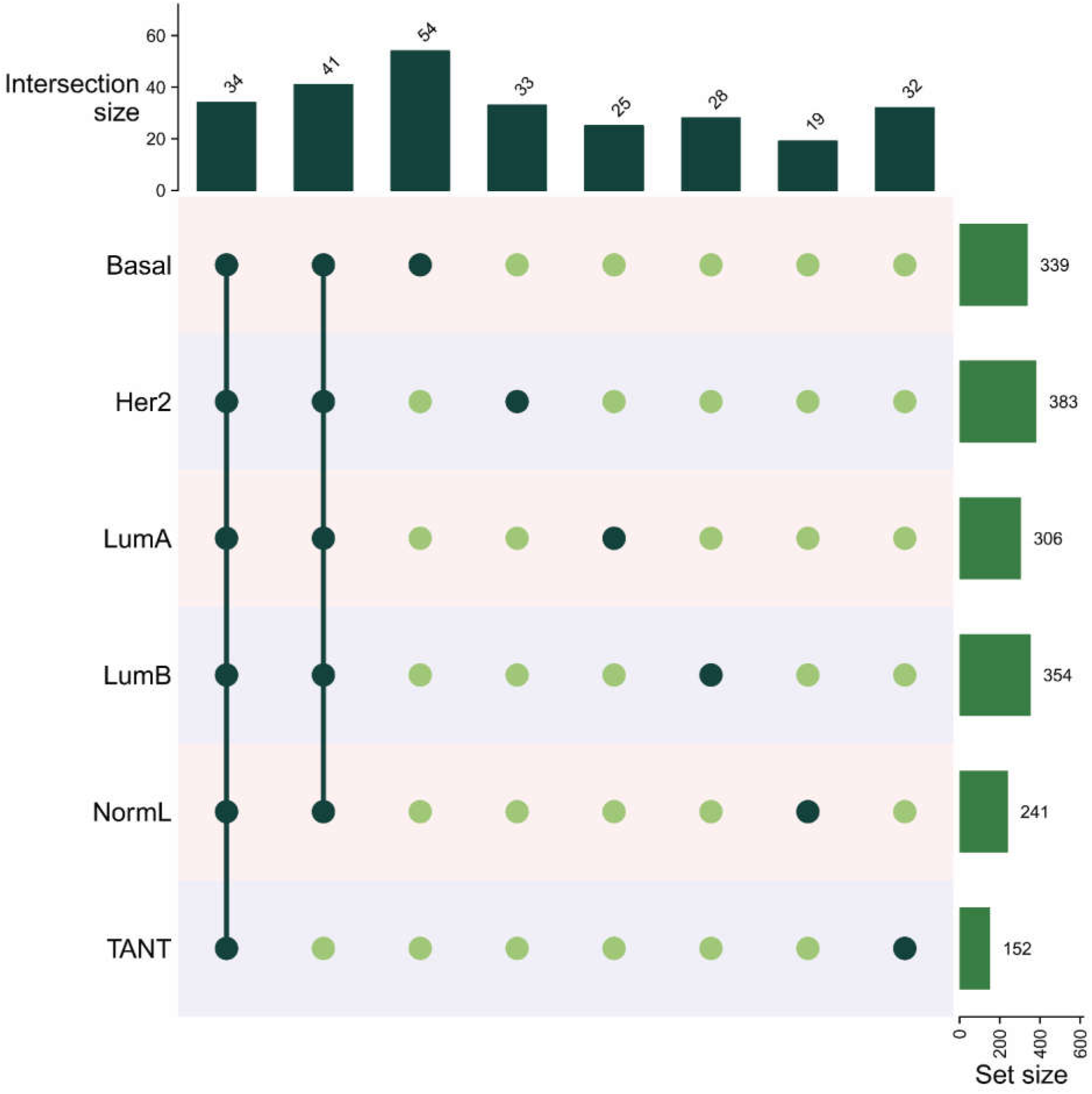
Distribution of common and unique hub proteins across all six tissue networks

**Figure 3.**
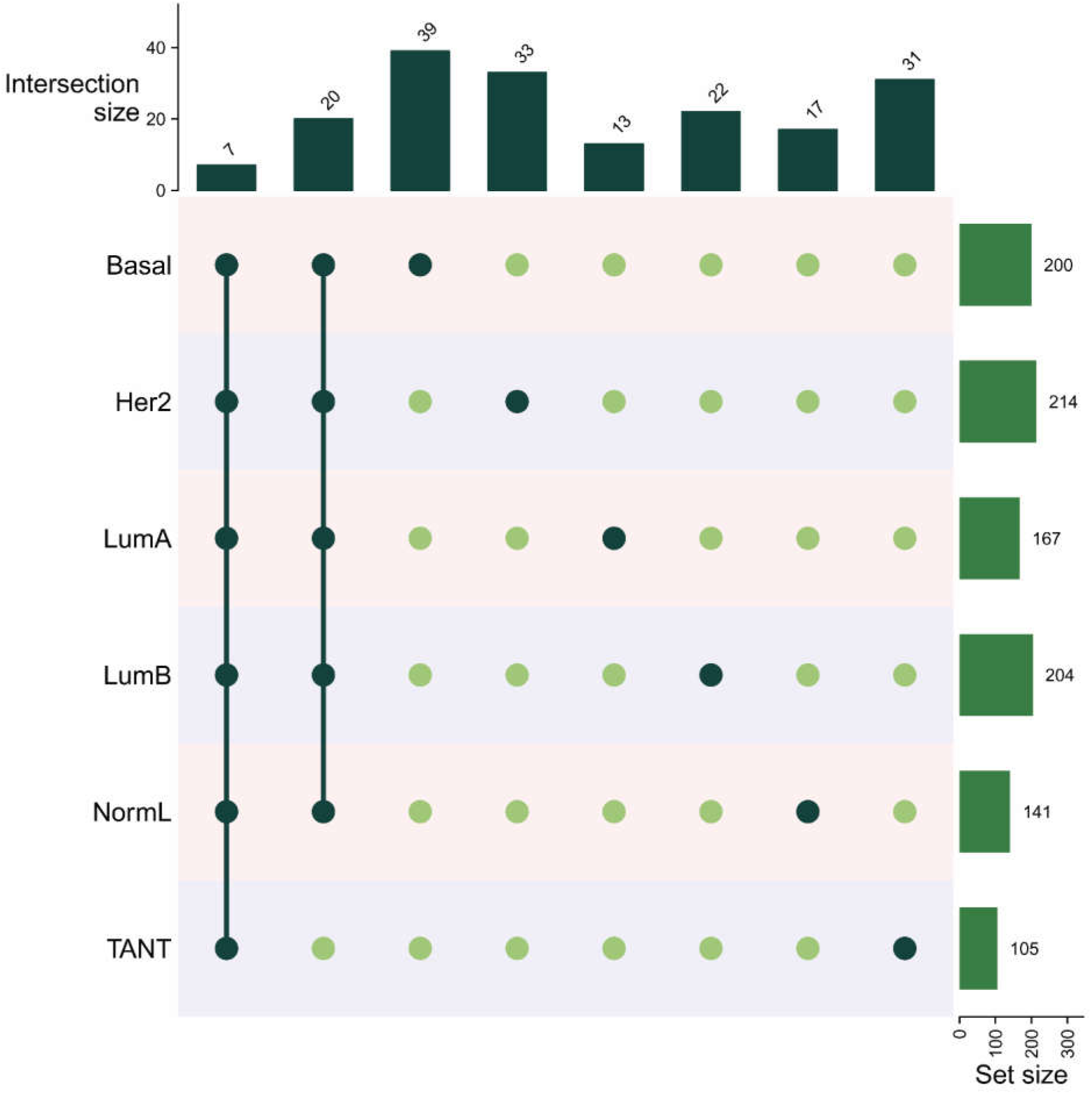
Distribution of common and unique bottleneck proteins across all six tissue networks

### Vulnerable proteins and protein pairs of tumor and TANT networks

Proteins with 14 calculated properties were processed to identify VPs for each tissue type. These VPs of tumor tissues and TANTs were identified with the help of boxplot-based criteria of detecting outliers in the following two steps: (i) proteins falling beyond the limit of either lower or upper whisker depending on the topological property as implemented in the NetVA package were considered outliers, and (ii) all outlier proteins, as per any three out of 14 topological properties, were considered as VPs. This procedure was repeated for each tissue type, and the list of corresponding VPs was identified. Further, VPPs for each tissue network were identified by deleting two VPs in pairs connected with each other from all VPs in the corresponding tissue network. This was followed by calculating all 14 topological properties of the resultant network. Subsequently, the VPPs were identified by following the same criteria as the VPs. Further, this procedure was repeated for each tissue type, and the corresponding list of VPPs was identified. It was observed that there were several VPs and VPPs that were present across all six tissue networks (Figs. 4 and 5). The detailed distribution of VPs across all six tissue networks is available in Figs. 4 and 6. Moreover, all the commonly present VPPs were found participating in two protein complexes across all six tissue networks (Fig. 7). The detailed lists VPs and VPPs across all six tissues are available in Tables S3 and S4.

**Figure 4.**
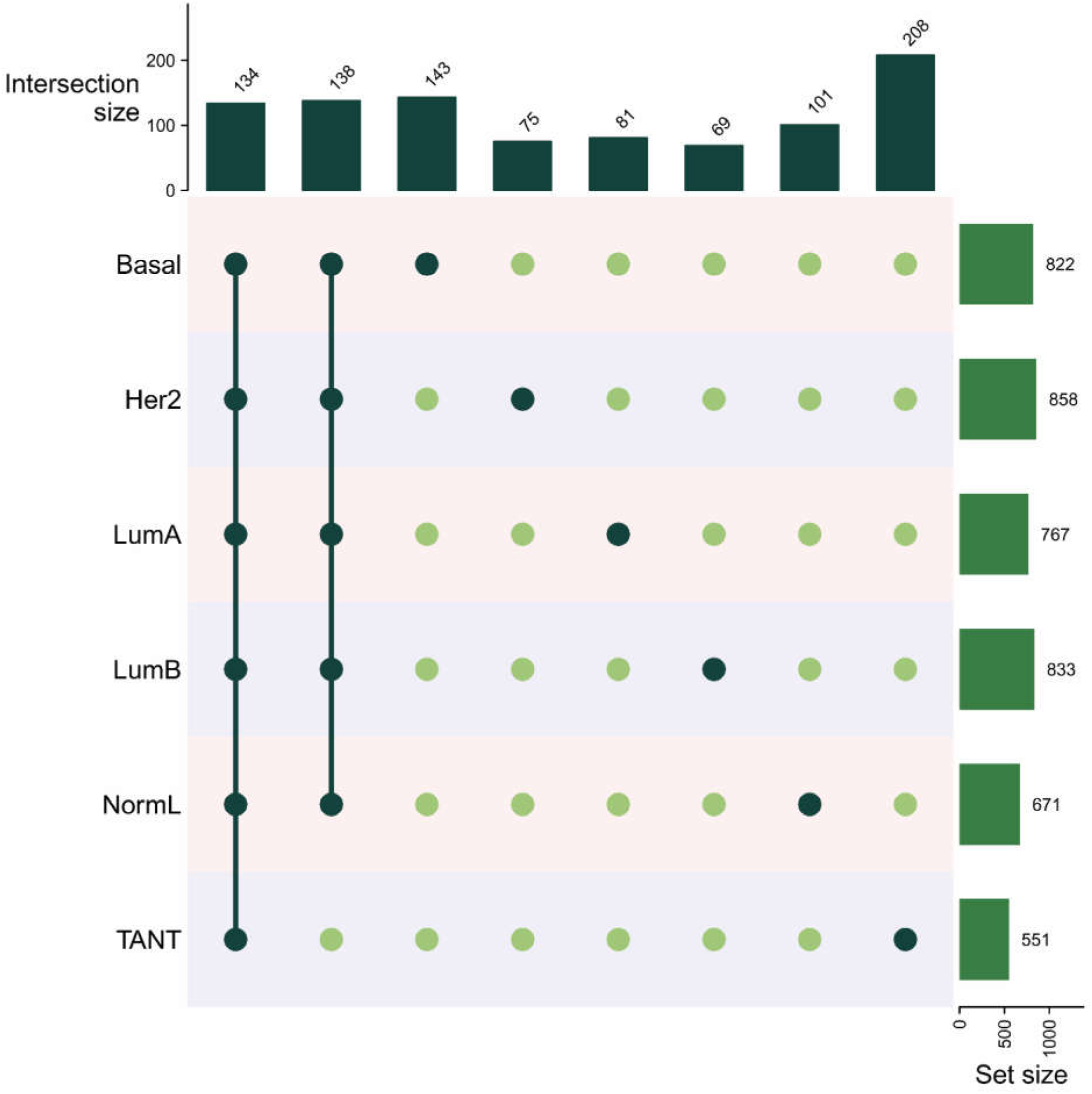
Distribution of common and unique VPs across all six tissue networks

**Figure 5.**
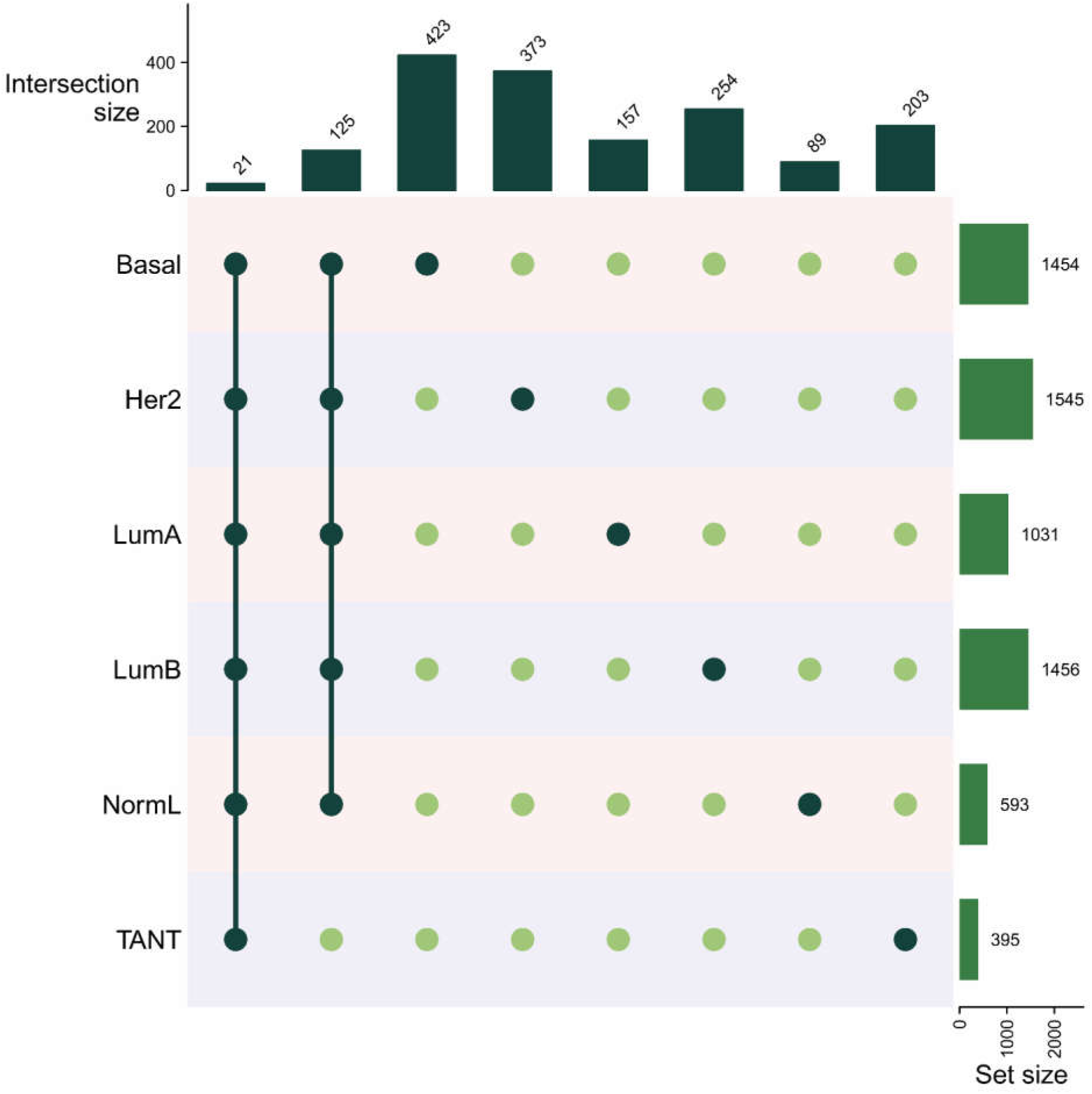
Distribution of common and unique VPPs across all six tissue networks

**Figure 6.**
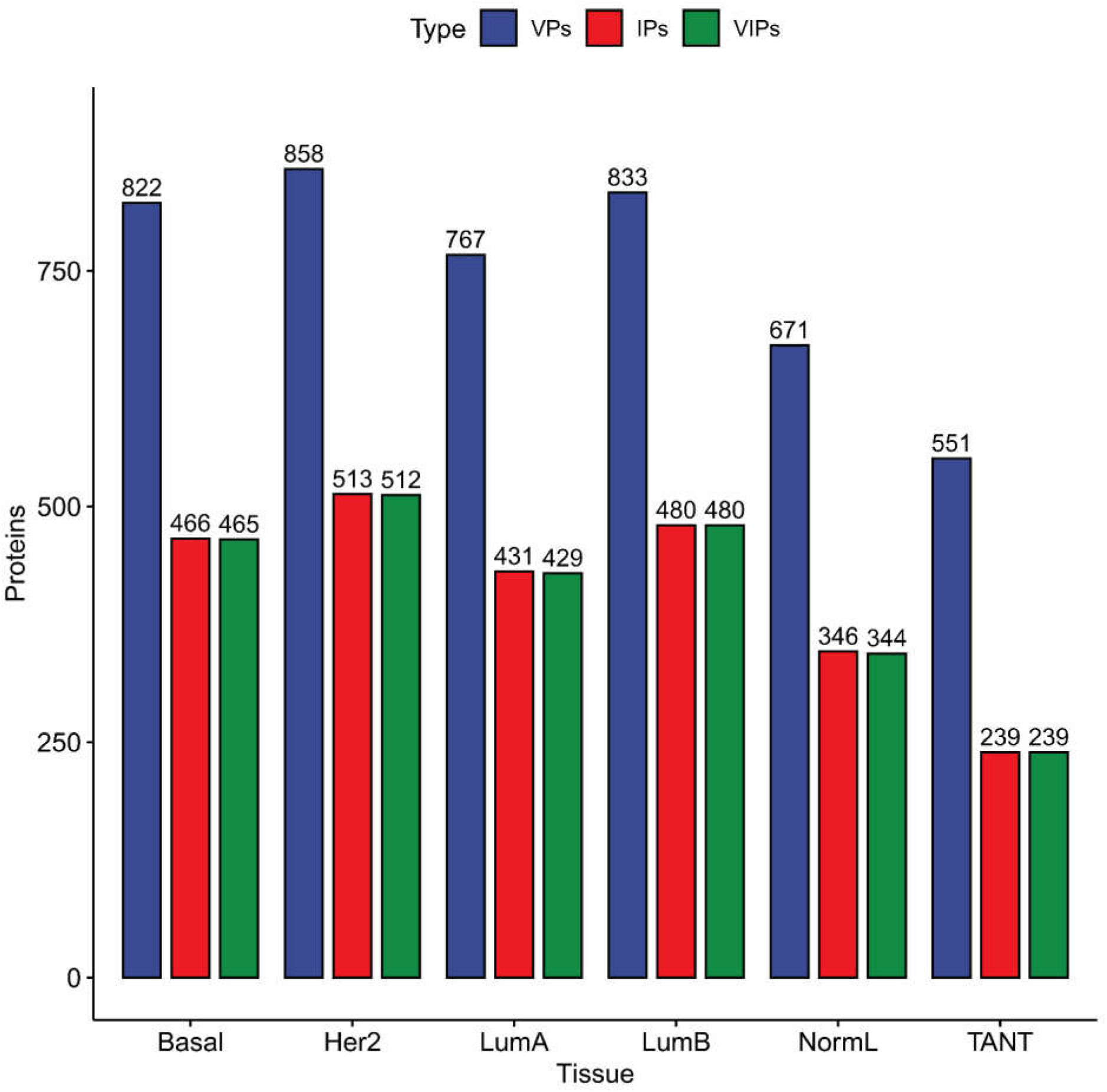
Distribution of VPs, IPs, and VIPs across all six tissue networks

**Figure 7.**
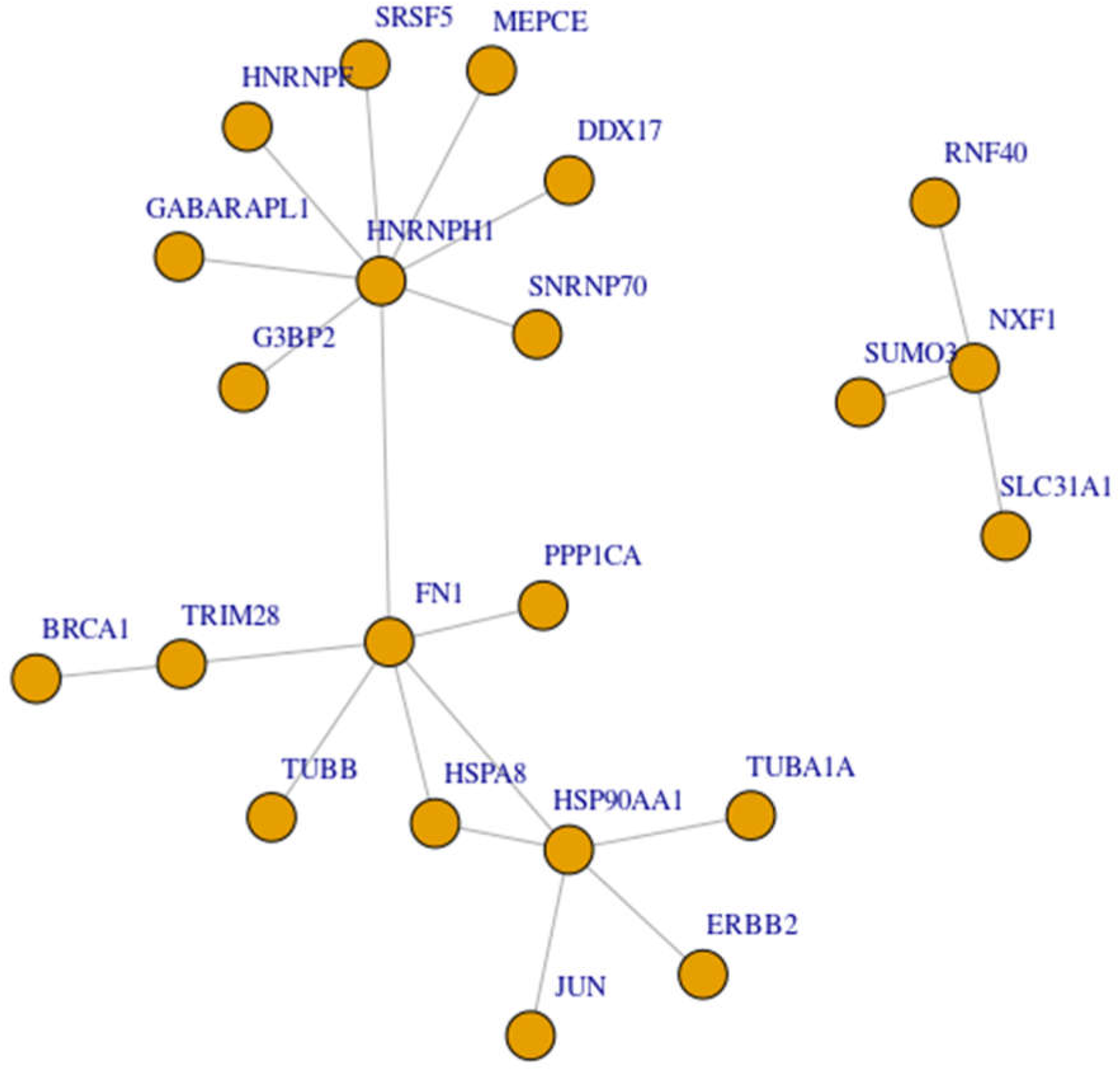
Common VPPs forming two common complexes composed of VPs and conserved across all six tissue networks

### Influential proteins of tumor and TANT networks

IPs for each tissue type were identified by analyzing the corresponding tissue networks using the “evc” function, as mentioned in the Methods section. Further, we also identified proteins that were vulnerable as well as influential (VIPs) in the case of each tissue network by checking their presence in the corresponding list of VPs. The distribution of IPs and VIPs in the case of each tissue type is available in Figs. 6 and 8. The detailed lists of IPs across all six tissues are available in Table S5.

**Figure 8.**
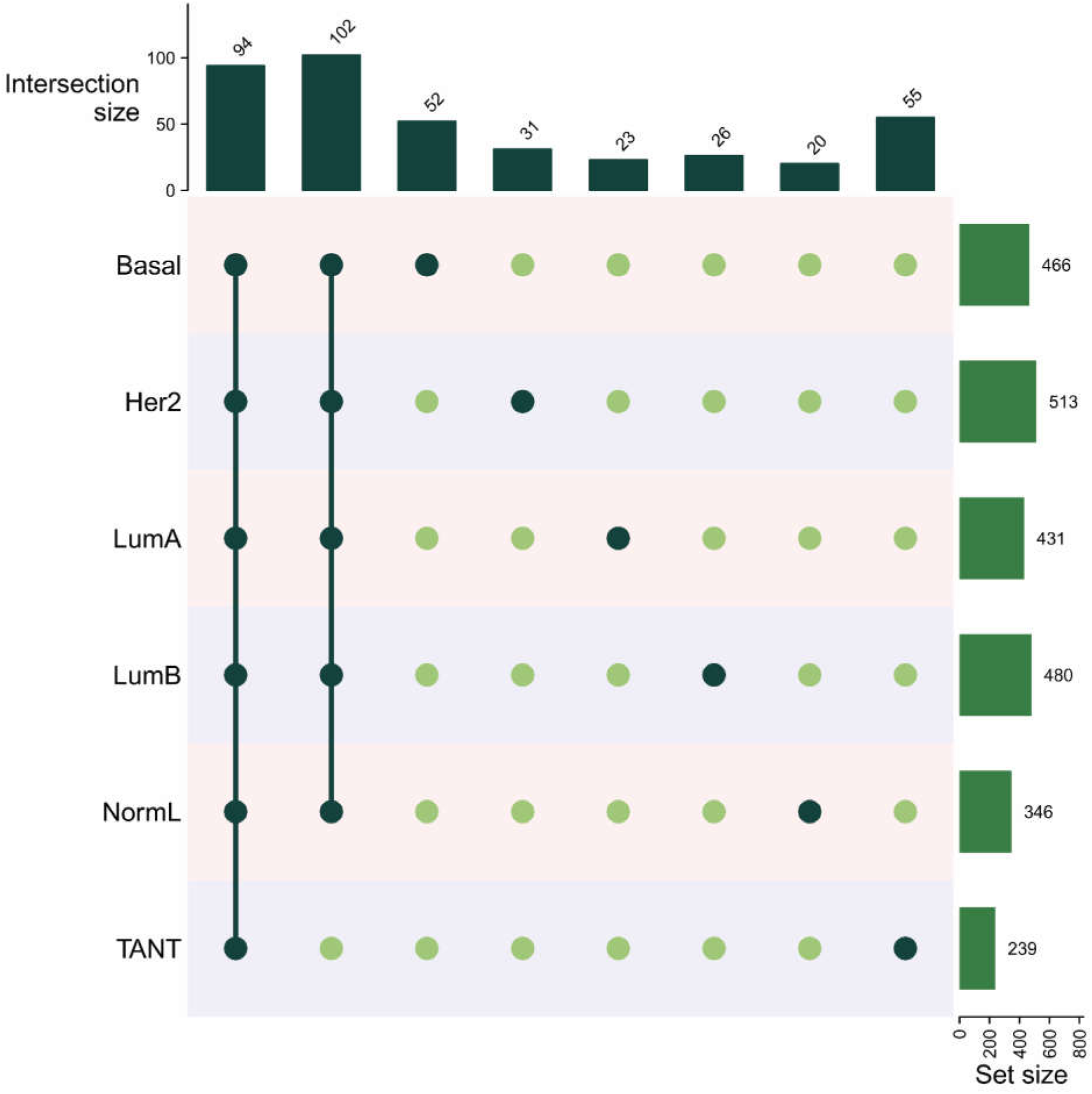
Distribution of common and unique IPs across all six tissue networks

### Vulnerable protein complexes of tumor and TANT networks

Potential protein complexes for each tissue type were predicted using the ClusterOne software. Subsequently, VPs were mapped onto these predicted protein complexes to identify VPCs. This procedure was repeated for each tissue type. As a result, it was found that there were no protein complexes that were entirely composed of VPs across all six tissue types. Consequently, all protein complexes with at least three VPs present therein were identified as VPCs for all six tissue networks. The detailed distributions of these VPCs across all six tissue networks are available in Table 2.

**Table 2.** Distribution of clusters and VPCs across all six tissue networks.

| Network | Clusters | VPCs |
| --- | --- | --- |
| Basal | 496 | 18 |
| Her2 | 497 | 15 |
| LumA | 413 | 13 |
| LumB | 492 | 14 |
| NormL | 331 | 8 |
| TANT | 219 | 4 |

VPCs, vulnerable protein complexes.

### Vulnerable, influential, hub, and bottleneck proteins associated with diseases

We checked the association of all key proteins (viz., VPs, IPs, hubs, and bottlenecks) with any diseases, including cancers, using the DisGeNET data on gene-disease associations. Only gene-disease associations reported in at least two published literature were considered for this. The detailed distribution of disease-associated VPs, IPs, hubs, and bottlenecks is available in Table 3.

**Table 3.** Distribution of disease-associated proteins among VPs, IPs, hubs, and bottlenecks.

| Tissue | VPs | IPs | Hubs | Bottlenecks |
| --- | --- | --- | --- | --- |
| Basal | 778 | 445 | 321 | 195 |
| Her2 | 803 | 489 | 364 | 205 |
| LumA | 721 | 416 | 294 | 160 |
| LumB | 779 | 459 | 340 | 196 |
| NormL | 639 | 333 | 235 | 136 |
| TANT | 517 | 233 | 147 | 101 |

## Discussion

Cancer studies involving patients’ tumor tissue samples generally consider TANTs as controls. However, it is well established that TANTs are not completely normal; instead, they possess various tumorigenic features, including mutated and over-expressed genes, among others [Aran et al., 2017; Kumar & Vindal, 2023]. Therefore, it is essential to explore the TANTs along with tumor tissues to study the tumorigenesis of cancers. It will help identify key proteins expressed in TANTs beside tumor tissues that will further aid in developing early-stage-specific and more efficient biomarkers and therapeutic candidates for better management and treatment of the disease. The prediction of key proteins that result in the increased vulnerability or reduced robustness of a particular system is of great importance because the vulnerable system has the capability of transforming normal healthy cells into diseased cells. These key proteins and the vulnerable system help to estimate the effect of targeted dysregulations or perturbations, such as drug-induced silencing or inactivation of the disease-associated physiological and pathological states. In this direction, the present study constructs and analyzes six PPI networks of different breast tissues, viz., Basal, Her2, LumA, LumB, NormL, and TANT, through network influence and vulnerability analysis.

Exploration of PPI networks of all six tissues using degree and betweenness led to identifying hubs and bottlenecks, respectively. Among all hubs, there were 34 proteins (viz., ADAMTSL4, APLNR, AQP3, AR, CD79A, DYNLL1, DYNLL2, ECT2, FBXO6, FN1, FOXA1, GABARAPL1, GAPDH, H2AX, HNRNPDL, HNRNPF, HSPA4, HSPA8, JUP, PAN2, PICK1, PPP1CA, RBM39, SNRNP70, SRSF5, SSX2IP, SUMO3, SUV39H1, TK1, TRIM28, TSC1, USHBP1, ZC3HAV1, and ZDHHC17) commonly present as hubs across all six tissue networks. Further, seven proteins (viz., ECT2, ERBB3, EZH2, GABARAPL1, HSPA1A, JUN, and ZDHHC17) were commonly present bottlenecks across all six tissue networks. Some proteins across all six tissue networks were hubs as well as bottlenecks, such as GABARAPL1, ECT2, and ZDHHC17.

Upon vulnerability analysis of all six tissue networks, we identified some key proteins as VPs, which have been previously reported to be involved in the tumorigenesis of breast cancer. There were 134 VPs, including some well-known cancer gene products, viz., AR, BRCA1, CCND1, ERBB2, ERBB3, JUN, FN1, FOXA1, NRAS, and STAT1, commonly present across all six tissue networks. It has been previously reported that androgen and AR have an involvement in breast cancer tumorigenesis and metastasis [Feng et al., 2017]. The gene encoding BRCA1, a tumor suppressor having binding activity associated with DNA damage response, has been reported in breast cancer as a mutated gene [Sjoblom et al. 2006]. Further, the overexpression of the gene encoding ERBB2 has been reported to crank up breast cancer metastasis [Tan et al. 1997; Holbro et al. 2003]. A study has also reported that the overexpression of the gene encoding JUN in breast cancer cells provides a more aggressive phenotype and, hence, causes breast cancer metastasis [Zhang et al. 2007]. Next, the gene encoding FOXA1 had an expression correlation with the LumA subtype and was a cancer-specific and significant predictor of survival in ER+ breast cancer patients [Badve et al. 2007].

Further, among VPs of each tissue network, some VPs were hubs or bottlenecks, whereas some were neither hubs nor bottlenecks. This revealed that most VPs of a network can not necessarily be only proteins occupying central positions in the network or proteins connected with a large number of other proteins present therein. When we analyzed these networks again by deleting VPs in pairs connected with each other, we could identify VPPs across all six tissue networks. Among the VPPs of each tissue network, there were 21 VPPs (viz., BRCA1-TRIM28, HSP90AA1-HSPA8, HSP90AA1-FN1, HSP90AA1-ERBB2, HSP90AA1-TUBA1A, HSP90AA1-JUN, DDX17-HNRNPH1, SRSF5-HNRNPH1, RNF40-NXF1, SNRNP70-HNRNPH1, HSPA8-FN1, FN1-TRIM28, FN1-HNRNPH1, FN1-PPP1CA, FN1-TUBB, SLC31A1-NXF1, G3BP2-HNRNPH1, GABARAPL1-HNRNPH1, MEPCE-HNRNPH1, NXF1-SUMO3, HNRNPH1-HNRNPF) commonly present across all six tissue networks. Herein, it can be observed that these VPPs were participating in the formation of one large complex with 18 interactions among 18 VPs and one small complex with three interactions among four VPs (Fig. 5). These two protein complexes were conserved across all six tissues, and their edges composed of VPPs were most vulnerable compared to others. Member proteins of the larger complex were mainly associated with gene-ontology (GO) annotations, such as sequence-specific DNA binding, RNA binding, protein binding, hydrolase activity, phosphoprotein phosphatase activity, GTP binding, and structural molecule activity. At the same time, member proteins of the smaller complex included GO annotations, such as protein homodimerization activity, ubiquitin-protein transferase activity, copper ion transmembrane transporter activity, and SUMO transferase activity, besides RNA binding and nucleotide binding.

When we identified the most influential proteins, i.e., IPs for each tissue type, based on their positions and interactions in the respective network, it was observed that almost all IPs were vulnerable, as present in the corresponding list of VPs across all six tissue networks. Similarly, it was noticed that all hubs and bottlenecks were vulnerable across all six tissue networks. However, most hubs and bottlenecks were influential, but not all across all six tissue networks. These observations were consistent with our previous study [Kumar et al., 2024]. Furthermore, we identified 94 proteins as shared IPs across all six tissue networks. These common IPs were found vulnerable as well and included several previously known cancer-associated proteins, such as AR, AURKA, BRCA1, ERBB2, ERBB3, FN1, FOXA1, JUN, MKI67, NRAS, STAT1, SUMO3, and TRIM28.

The exploration of all key proteins (viz., VPs, IPs, hubs, and bottlenecks) for their association with diseases, including cancers, indicated that the majority of them were previously known for their association with various diseases as per the gene-disease association data available in the DisGeNET. Besides these known disease-associated gene-encoding proteins, a few proteins have not been reported in the case of any disease. This proves the potential of the current study and the methods adopted herein in efficiently identifying a list of key candidate proteins, including previously known and novel ones. These key proteins, especially TANT, can be utilized further as putative biomarkers and/or therapeutic candidates to devise better management of the disease at an early stage.

## Conclusions

We explored the protein interaction networks of TANTs and five breast cancer subtypes using network vulnerability analysis implemented in the NetVA. This, along with the detection of hubs, bottlenecks, and influential proteins, helped us to identify some key proteins and protein pairs that participate in the formation of protein complexes and have distinct and conserved patterns of organization across all six tissue networks. Further, the presence of novel proteins without any known association with diseases, along with known proteins having a well-documented association with diseases, including cancers, highlighted the study’s efficacy in identifying key proteins associated with the tumorigenesis of breast cancer. However, these candidate molecules, especially common VPs across all six tissue networks, must be explicated further using other experimental approaches, such as *in vitro* and *in vivo,* to devise more effective diagnostic and therapeutic strategies.

## Acknowledgments

VV would like to thank the Indian Council of Medical Research (ICMR), New Delhi (ISRM/12(72)/2020, ID: 2020-2951), and the Institution of Eminence (IoE) – University of Hyderabad (No. UoH/IoE/RC3-21-052) for their financial support. SK also acknowledges the IoE - University of Hyderabad for the performance-based publication incentives and the ICMR for the Senior Research Fellowship (Grant No.: 3/2/2/113/2019/NCD-III, ID: 2019-6723). Further, the authors thank the Center for Modeling, Simulation & Design (CMSD), University of Hyderabad for providing computational facilities.

## Author Contributions

SK and VV conceived and designed the study. SK and AA curated and filtered the PPI data from various sources. SK performed the experiments and analyzed the data. SK drafted, wrote, and edited the manuscript. VV reviewed and edited the manuscript. VV supervised the study. All authors read and approved the final manuscript.

